## Supplementary figures and images for "Transplantation of exogenous mitochondria mitigates myocardial dysfunction after cardiac arrest"

### supplemental Fig. 1

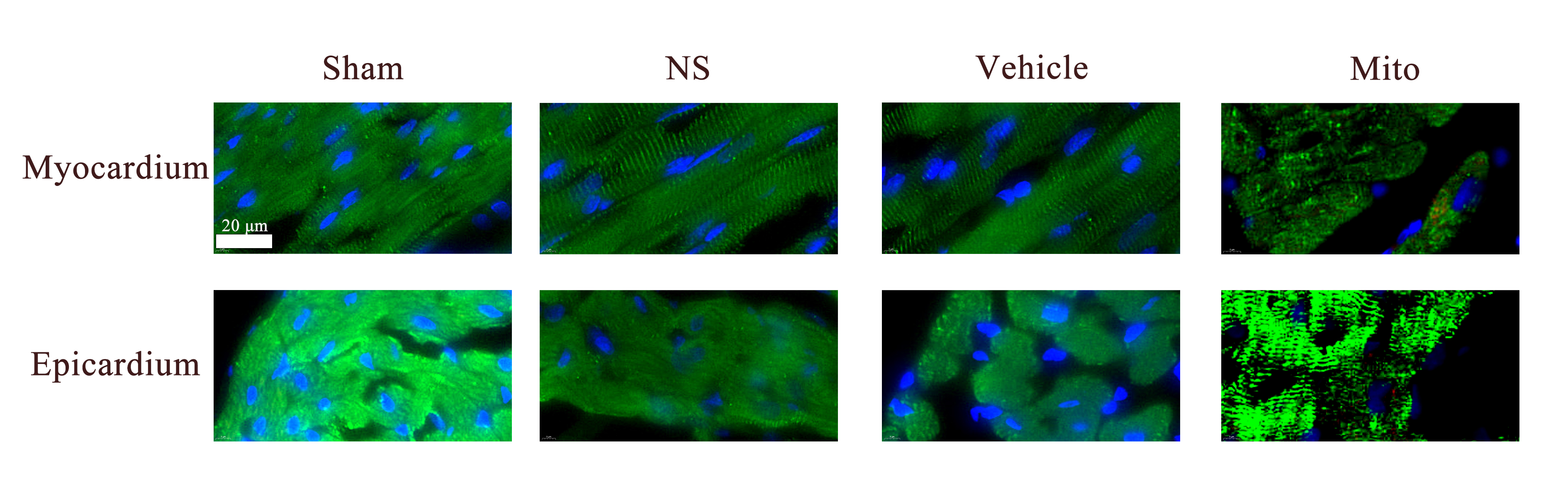
