## Supplemental Table S1 for "Transplantation of exogenous mitochondria mitigates myocardial dysfunction after cardiac arrest"

**Table S1 Details of antibodies used in the methodology**

| **Variable** | **Manufacturer** | **Catalog number** | **Dilution** | **Validation** |
| --- | --- | --- | --- | --- |
| **COX- IV** | Beyotime | AC610 | 1:1,000 | WB |
| **Caspase-3** | Cell Signaling Technology | 14220T | 1:1000 | WB |
| **GAPDH** | Epizyme | LF206 | 1:3,000 | WB |
| **Horseradish peroxidase-labeled secondary antibody** | Beyotime | A0208 | 1:2000 | WB |
| **Alpha-actinin 2** | GeneTex | GTX103219 | 1:500 | IF |
| **Alexa Fluor 488-labeled Goat anti-Rabbit IgG** | Huzhen | HZ0176 | 1:1,000 | IF |
| **DAPI** | Beyotime | P0131-5 mL | No dilution | IF |
| **Tom20 antibody** | Affinity | AF5206 | 1:1000 (WB) 1:100 (IF) | WB, IF |
| **Anti-Rabbit IgG (H+L) Antibody, Peroxidase-Labeled** | SeraCare | 5220-0336 | 1:400 | IF |

WB, western blot; IF, immunofluorescence.
