## Supplemental Table S2 for "Transplantation of exogenous mitochondria mitigates myocardial dysfunction after cardiac arrest"

Table S2 Scoring Standard of Myocardial Pathological Injury

| Grade | Scoring Standard | Score |
| --- | --- | --- |
| 0 | The staining of myocardial fiber was homogeneous, cross striation was distinct, myocardial interstitium has no sign of inflammatory cellular invasion, no sign of hemorrhagic necrosis; | 0 |
| Ⅰ | Partial myocardial fiber presents wave shape, myoplasm distribution was inhomogeneous, myocardial interstitium has no sign of hemorrhage; | 2 |
| Ⅱ | Partial myocardial fiber breakage is noted, with focal hemorrhage, myocardial interstitium has inflammatory cellular hemorrhage; | 4 |
| Ⅲ | Partial myocardial has focal necrosis, myocardial interstitium has hemorrhage and inflammatory cellular invasion; | 6 |
| Ⅳ | A majority of myocardium has focal necrosis, myocardial interstitium has hemorrhage and inflammatory cellular invasion; | 8 |
| Ⅴ | A majority of myocardium has focal necrosis, myocardial interstitium has diffusive hemorrhage and inflammatory cellular invasion. | 10 |
